# Astrocyte Ca^2+^ signalling mediates long-distance metaplasticity in the hippocampal CA1

**DOI:** 10.1101/2023.06.19.545623

**Authors:** Owen D. Jones, Anurag Singh, Barbara J. Logan, Wickliffe C. Abraham

## Abstract

Astrocytes play an increasingly recognised role in regulating synaptic plasticity, but their contribution to metaplasticity is poorly understood. We have previously described a long-distance form of metaplasticity whereby priming stimulation in stratum oriens inhibits subsequent LTP in the neighbouring stratum radiatum of the hippocampal CA1 region of both rats and mice. Using genetic and pharmacological strategies to manipulate astrocytic Ca^2+^ signalling, we now show this form of metaplasticity requires inositol triphosphate receptor-dependent Ca^2+^ release in these cells. Blocking Ca^2+^signalling or inositol triphosphate receptors in single radiatum astrocytes abolishes the metaplasticity at nearby synapses. We also show the relevant Ca^2+^release in astrocytes is driven by adenosine A_2B_ receptors, and stimulation of these receptors elicits the metaplasticity effect both *in vitro* and *in vivo*. Further, the metaplasticity requires signalling via tumor necrosis factor, but this cytokine is required to act on astrocytes, not neurons. Instead, glutamate, acting on GluN2B-containing NMDA receptors, is the likely gliotransmitter that signals to neurons to inhibit LTP. Together these data reveal a novel role for astrocytes in hippocampal LTP regulation across broader spatiotemporal scales than previously recognised.

**Main points:**

- In hippocampal CA1, “priming” activity inhibits subsequent LTP at synapses hundreds of microns away.
- This effect requires astrocytic Ca^2+^signaling, and a molecular cascade involving adenosine A_2B_ receptors, tumor necrosis factor and GluN2B-containing NMDA receptors.
- The metaplasticity effect is evident *in vitro* and *in vivo*.
- Long-distance astrocyte signaling is a mechanism for regulating neural activity over broad spatiotemporal scales.

## 1. Introduction

Astrocytes regulate various forms of synaptic plasticity (Haydon & Nedergaard, 2015; Min, Santello, & Nevian, 2012), and are well-placed to exert ongoing regulation of plasticity (metaplasticity: Hulme, Jones, & Abraham, 2013) over varying spatiotemporal scales (Jones, 2015). However, with few exceptions (Panatier et al., 2006), the contribution by astrocytes to metaplasticity has not been investigated.

Previously we described a unique Bienenstock, Cooper and Munro (BCM)-like form of metaplasticity in hippocampal CA1, whereby “priming” stimulation delivered to afferents in the basal dendritic zone in stratum oriens (SO) inhibits subsequent long-term potentiation (LTP) hundreds of microns away in the apical dendritic zone in stratum radiatum (SR) (Hulme, Jones, Ireland, & Abraham, 2012). This long-distance metaplasticity effect occurs independently of canonical neuronal mechanisms such as cell-firing or N-methyl-D-aspartate receptor (NMDAR) activation during the priming stimulation. Instead it requires adenosine A2B receptor (A_2B_R) activation and functional connexin43 proteins (Jones, Hulme, & Abraham, 2013), the former enriched and the latter exclusively located on astrocytes (Boison, Chen, & Fredholm, 2010; Giaume et al., 1991; van Calker & Biber, 2005). Thus, in contrast to the cell-autonomous BCM model (Bienenstock, Cooper, & Munro, 1982), communication with astrocytes could be critical for this long-range metaplasticity effect. Supporting this possibility, priming stimulation in SO triggers Ca^2+^elevations in astrocytes both locally in SO and distally in SR (Hulme, Jones, Raymond, Sah, & Abraham, 2014). However, a direct role for astrocyte Ca^2+^ signalling has not yet been established for any of the priming protocols that we have used to date. Further, while we have shown that the cytokine tumor necrosis factor (TNF) is also critical to the metaplastic state (Singh, Jones, Mockett, Ohline, & Abraham, 2019), we have not previously established a connection between TNF or adenosine and astrocyte Ca^2+^ in our model. Here, we demonstrate that blocking the calcium-dependent functionality of astrocytes in SR completely prevents the metaplasticity effect at nearby synapses, whether generated electrically or via administration of TNF or an A_2B_R agonist. We also show that inositol triphosphate (IP_3_)-gated stores in astrocytes are the relevant source of Ca^2+^. Finally, we demonstrate that the post-priming activation of GluN2B subunit-containing NMDA receptors mediates the LTP inhibition.

## 2. Materials and Methods

### 2.1 Animals

All experiments were conducted with the approval of the University of Otago Animal Ethics Committee, and in accordance with New Zealand legislation. Animals were obtained from colonies maintained at the University of Otago Breeding Station.

### 2.2 Drugs and reagents

Heparin and TNF were purchased from R&D Systems. BAY-60 6583, MRS1754, D-APV and MCPG were sourced from Tocris. Ifenprodil was purchased from Hello Bio. All other dyes and active compounds were sourced from Sigma. Stocks of BAY-60 6583 and MRS1754 were dissolved in DMSO. Stock TNF was suspended in PBS with 0.1% bovine serum albumin as a carrier protein. Stock MCPG was dissolved in 0.1 M NaOH. All stock solutions were diluted 1000x in aCSF for final working concentrations. Vehicle was delivered to slices in place of drug as a control where required.

### 2.3 In vitro experiments

#### Slice preparation

Acute, transverse hippocampal slices were prepared as previously described (Jones et al., 2013). Briefly, male rats or mice of either sex (6-8 weeks) were deeply anesthetised with 100 mg/kg i.p. ketamine (rat) or isoflurane inhalation (mouse) and decapitated. Brains were rapidly removed and placed in ice-cold dissection solution (mM: 210 sucrose, 26 NaHCO_3_, 2.5 KCl, 1.25 NaH_2_PO_4_, 0.5 CaCl_2_, 3 MgCl_2_, 20 D-glucose) gassed with 95% O_2_/5% CO_2_. For rat slices, hippocampi were dissected from Sprague-Dawley rat brains and area CA3 removed via knife cut. Hippocampi were then sectioned (400 µm) using a vibratome (Leica VT1000). For mouse slices, horizontal brain sections (400 µm) containing hippocampus cortex were prepared from each hemisphere. Slices were left to recover in a humidified chamber at the interface between air and extracellular aCSF solution (mM: 124 NaCl, 3.2 KCl, 1.25 NaH_2_PO_4_, 26 NaHCO_3_, 2.5 CaCl_2_, 1.3 MgCl_2_, 10 D-glucose) and gassed with 95% O_2_/5% CO_2_. Slices remained at 32°C for 30 min post-sectioning, then at room temperature for at least 90 min prior to transfer to the recording chamber. In patching experiments, sulforhodamine 101 (SR101, 0.5 µM) was included in the initial period of heated recovery to selectively label astrocytes (Nimmerjahn, Kirchhoff, Kerr, & Helmchen, 2004). Slices were then transferred to standard aCSF at room temperature.

#### Field recordings

For recordings, slices were transferred to a recording chamber and submerged in aCSF (32.5°C) at a flow rate of 2 ml/min. Synaptic field excitatory postsynaptic potentials (fEPSP, 2-3 mV amplitude at 65 μA) were evoked through separate stimulation of Schaffer collateral afferents in the SR and SO of hippocampal CA1 using 50 μm Teflon-insulated tungsten monopolar electrodes (diphasic pulse half-wave duration 0.1 ms). Electrodes were placed centrally within each stratum (midway between the stratum pyramidale (SP) and alveus for SO stimulation, and midway between the SP and stratum lacunosum-moleculare for SR stimulation). Recordings were made using glass micropipettes filled with aCSF (2-3.5 MΩ), placed approximately 400 μm from the stimulating electrode in the same stratum. Baseline stimulation consisted of single pulses delivered every 30 s, alternating between stimulating electrodes. Electrical priming stimulation in SO consisted of 3 × 100 Hz, 1 s trains spaced by 20 s, repeated after 5 min (Jones et al., 2013). In mouse experiments, this protocol was truncated to 2 x trains of TBS (i.e. 10 x 5 Hz bursts of 4 x 100 Hz pulses, repeated after 30 s). For LTP induction in SR, 10 × 5 Hz bursts of 4 x 100 Hz pulses (repeated after 30 s) were delivered 15-30 min after priming. LTP was assessed at 50-60 minutes post-induction, and expressed as the averaged percent of the baseline fEPSP slope. For glutamatergic blockade experiments, priming occurred in the presence of the AMPA/kainate receptor antagonist kyenurate (3 mM), the NMDAR antagonist D-APV (50 µM) and the metabotropic glutamate receptor (mGluR) antagonist MCPG (250 µM). This cocktail was bath-applied 15 min prior to and for the duration of priming. In ifenprodil experiments, the drug (3 µM) was present for the entire experiment. Inositol triphosphate receptor 2 knock out (IP3R2KO) mice were maintained as heterozygote pairs, yielding WT and KO littermate offspring for experimentation. Genotyping was performed as per the developers’ original publication (X. Li, Zima, Sheikh, Blatter, & Chen, 2005). The investigator was blind to genotype during data collection and analysis.

#### Astrocyte patching

Slices were submerged in aCSF (32.5°C) at a flow rate of 2 ml/min and imaged using differential interference contrast under a 40X objective attached to an upright microscope (Olympus BX50). SR101-positive cells in SR were identified in epifluorescence mode via green LED excitation (540 nm broadband, Mightex Corporation) and a long-pass orange-red filter (Omega Optical set XF34). SR101-positive cells were patched with pipettes (4-8 MΩ) containing standard intracellular solution (mM: KCH_3_O_3_S 135, HEPES 10, disodium phosphocreatine 10, MgCl_2_4, Na_2_ATP 4, NaGTP 0.4 (pH adjusted to 7.25 with KOH, 290–295 mOsM). Cells were confirmed as astrocytes if V_m_ was greater than -80 mV at rest and current-voltage tests revealed linear changes in V_m_ in response to depolarising current steps (200 pA) and no active membrane potentials. In various experiments, this solution was supplemented either with EGTA (0.45 mM) and CaCl_2_ (0.14 mM) to ‘clamp’ cytosolic Ca^2+^ levels at resting values (Henneberger, Papouin, Oliet, & Rusakov, 2010), or with heparin (1.5 mg/ml) to block IP_3_receptors (Power & Sah, 2007). In Ca^2+^ clamp experiments, the extracellular solution was supplemented with D-serine (10 µM) in all cases. Stimulating electrodes were placed ∼300 µm from the recording (patch) electrode and used to elicit synaptic activity. The resultant field synaptic potential, although attenuated, was readily detected via the patch recording electrode due to the weak resistance (< 5 MΩ here) of the astrocytic membrane (Henneberger & Rusakov, 2012). These astrocyte-detected field excitatory postsynaptic potentials (AfEPSPs) provided a stable measure of synaptic activity localised to the vicinity of the patched astrocyte.

Stimulating and recording parameters were conducted as per extracellular recordings, albeit on an abbreviated timescale. After a 15 min baseline, slices received either standard electrical priming in SO, or chemical priming via 10 min bath application of BAY 60-6583 (100 nM) or TNF (1.18 nM). After a further washout/recovery period (15-20 min), LTP was induced as described above and followed for 30 min.

### 2.4 In vivo experiments

#### Surgery

Adult male Sprague–Dawley rats (450–650 g) were anesthetized with isoflurane, placed in a stereotaxic frame, and maintained at 37 ± 0.3°C using a heat pad. Using standard surgical procedures, a 23 ga stainless-steel guide cannula was lowered into the right lateral ventricle for drug delivery (−0.6 mm posterior and 1.5 mm lateral to bregma, and 3.4 mm in depth from dura). Stimulating and recording electrodes (75 μm, Teflon coated stainless steel) were then implanted in the right hemisphere to establish Schaffer collateral-evoked field potentials recorded in the stratum radiatum of CA1. Using electrophysiological guidance, recording electrodes were positioned 3.3 mm posterior and 1.8 mm lateral from bregma. Stimulating electrodes were placed 3.8 mm posterior and 2.0 mm lateral to bregma. The guide cannula and electrodes were secured in place using dental cement, and positioning was confirmed histologically post-mortem. The initial slope of the negative-going fEPSP was used as a measure of synaptic efficacy. Following surgery, animals were housed individually and exposed to a normal 12 h light-dark cycle and given pain relief via carprofen for < 5 days.

#### Electrophysiology

Beginning 2 weeks post-surgery, animals were taken to a recording room and tested for usable field potential recordings evoked by Schaffer collateral stimulation (fEPSP slope 0.5–2 mV/ms at 200 μA). Baseline testing (0.05–0.017 Hz, 150 µs pulses) was undertaken for 30 minutes using a stimulus strength that elicited a fEPSP of 50% maximum slope. Baseline recordings were made 2–3 times per week at the same time during the animal’s light cycle until a stable level of evoked responses (varying by less than ± 10%) was obtained for at least 4 consecutive sessions. On the day of LTP induction, a 20 min period of baseline recording was undertaken before an experimental solution (15 µM BAY-60 6583 and/or 150 µM MRS1754 in 0.1% DMSO or PBS/DMSO control vehicle) was injected into the ipsilateral lateral ventricle with a 30 ga cannula protruding 0.5 mm below the guide cannula, using a Harvard Apparatus pump (5 µl at 1.0 µl/min). Test pulses then resumed for 30 min prior to LTP induction via TBS (4 trains spaced 30 s apart, each train being 10 x 5 Hz bursts of 5 pulses at 200 Hz; stimulation at 150% of test pulse intensity). Responses were followed for 60 min post-HFS and LTP was calculated as the average percent change in fEPSP slope from baseline at 50-60 min post-induction.

### 2.5 Data analysis

In extracellular experiments, *in vitro* and *in vivo* data were acquired and analysed using custom designed software based on the LabVIEW library. In patching experiments, data were acquired and analysed using pClamp version 10 (Molecular Devices). The initial slopes of fEPSPs and AfEPSPs were measured off-line and expressed as a percentage of baseline value (the averaged fEPSP slope for the 10 min preceding LTP induction). All data are expressed as mean ± SEM. Statistical comparisons were performed using Prism 9 software via one-way ANOVA or Student’s *t* test (paired or unpaired) as appropriate. A *p* value of < 0.05 was considered significant. For multiple comparisons, post-hoc testing was performed via Fisher’s LSD post-hoc test. Thus, in figures containing multiple comparisons, figure captions contain main ANOVA results and significant post-hoc differences are denoted via asterisks (*, *p* < 0.05; **; *p* < 0.01; ***, *p* < 0.001).

## 3. Results

### 3.1 Long-distance hippocampal metaplasticity requires astrocytic Ca^2+^ stores

We first bath-applied a cocktail of glutamatergic antagonists (kyenurate, APV and MCPG) to adult rat hippocampal slices. This cocktail not only blocked NMDARs and mGluRs during priming as in our previous report (Hulme et al., 2012), but also blocked AMPARs and KARs. Thus, glutamatergic transmission, and therefore postsynaptic depolarisation and feed-forward GABAergic signalling, were effectively blocked during SO priming. Despite this treatment, LTP induced 30 min after drug washout in primed slices was significantly inhibited compared to that in drug-treated but non-primed slices (**Fig. 1a**). These data confirm that the metaplasticity effect is not due to glutamatergic or intraneuronal voltage-based communication between SO and SR synapses, or recruitment of GABAergic interneurons. Accordingly, we sought to find evidence for a non-neuronal mechanism for mediating the long-range signalling between CA1 strata, such as that played by the astrocytic network.

**Figure 1:**
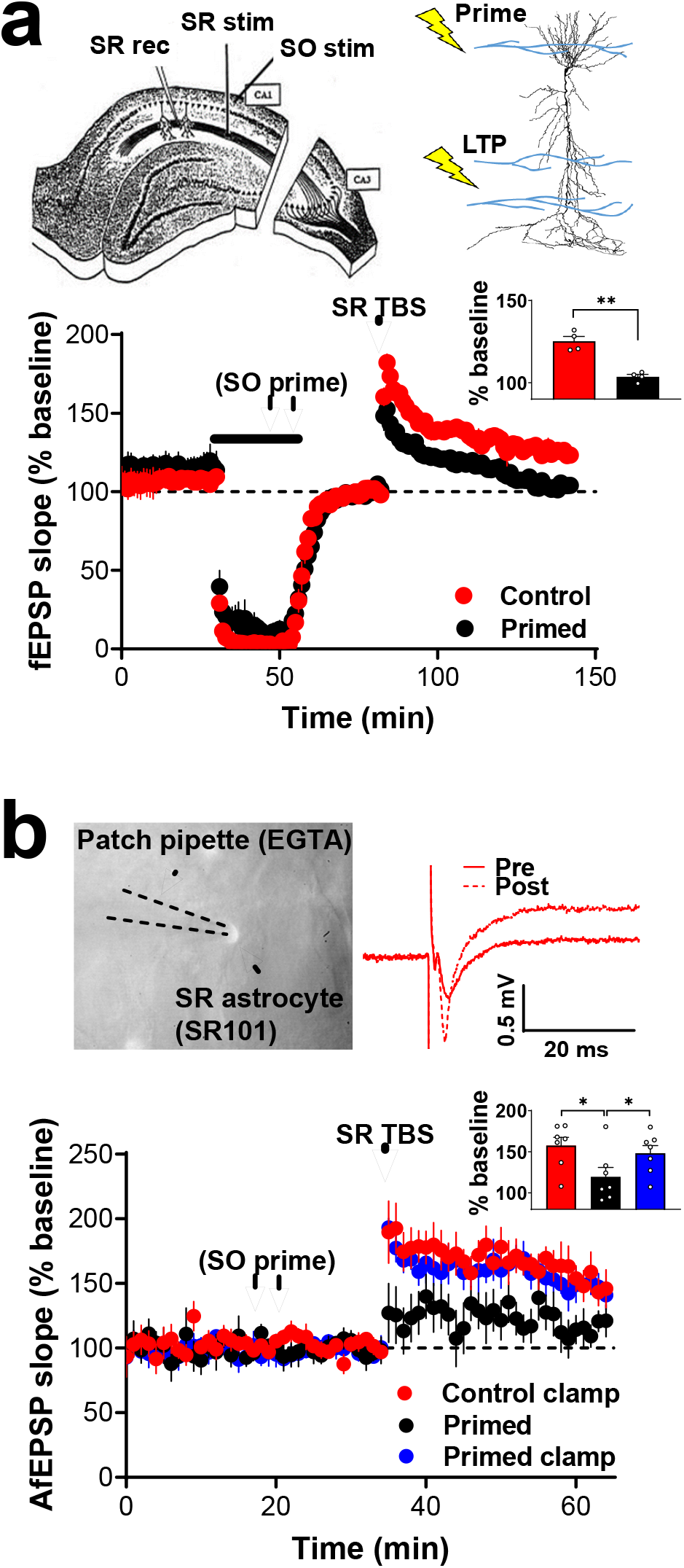
Long-distance hippocampal metaplasticity requires astrocytic Ca^2+^. **a**, Upper: Electrode placement in slices, with stimulation in SO (priming) and SR (LTP). Lower: Glutamatergic blockade via kyenurate, APV and MCPG (black bar) did not alter the inhibitory effects of SO priming on subsequent SR LTP (Control (n=4): 125 ± 3%; Primed (n=4): 104 ± 1%; *t*_(6)_ = 2.45, *p* < 0.001). **, *p* < 0.01. Inset: Bar chart of average LTP during the final 10 min of recording. **b**, Upper: EGTA-filled patch pipette attached to an SR101-labelled astrocyte in SR, and representative field potentials recorded through the astrocyte membrane (AfEPSP) before (pre) and after (post) LTP induction. Lower: Ca^2+^ clamp of astrocytes in SR blocked the metaplastic effects of SO priming (Control + Clamp (n=7): 158 ± 10%; Primed (n=7): 119 ± 12%; Primed + Clamp (n=7): 148 ± 9%; *F*_(2,18)_ = 3.67, *p* = 0.046). Inset: Bar chart of average AfEPSP LTP during the final 10 min of recording.

Following established methods (Henneberger & Rusakov, 2012), we recorded field excitatory postsynaptic potentials (AfEPSPs) in SR through pipettes attached to single whole-cell patch-clamped SR astrocytes, and used internally delivered EGTA (Henneberger et al., 2010) in some cells to “clamp” astrocyte Ca^2+^at basal levels (with 10 μm D-serine present in the artificial cerebrospinal fluid (aCSF) to enable LTP induction). The priming effect was still evident when recording fEPSPs in this way through the patch electrode (**Fig. 1b**). In striking contrast, recording under Ca^2+^clamp conditions caused the locally recorded priming effect to be blocked, restoring LTP induction to control levels (**Fig. 1b**). In separate control experiments, we verified the actions of the clamp on astrocytic Ca^2+^by testing LTP induction in aCSF without D-serine. Under these conditions, LTP could not be induced locally to the clamped astrocyte (**Fig. S1**), validating the effectiveness of this protocol as per its initial report (Henneberger et al., 2010).

### 3.2 Inositol triphosphate-gated stores are the relevant source of astrocytic Ca^2+^

The next question we addressed was the source of the astrocytic calcium needed to generate the metaplasticity effect. We have previously demonstrated a role for inositol triphosphate (IP_3_)-gated Ca^2+^stores in generating the metaplasticity effect (Hulme et al., 2012), but without knowing the cell type involved. Accordingly, we tested whether these critical stores are located in astrocytes. When the IP_3_ receptor blocker heparin was included in the astrocyte patch pipette solution, it did not affect control LTP in non-primed slices, as expected, but it completely blocked the priming-mediated inhibition of LTP (**Fig. 2a)**. To confirm these results, we turned to IP3R2KO mice, which lack the astrocyte-specific IP_3_ receptor subtype 2 and display diminished Ca^2+^release from astrocytic G-protein coupled stores but display normal LTP (Agulhon, Fiacco, & McCarthy, 2010). Here we found that while the metaplasticity effect was evident in wild-type mice, priming had no effect on later LTP in slices from littermate knock-out mice (**Fig. 2b**). Thus, IP_3_ R-gated release of calcium from intracellular stores in astrocytes located in the SR plays a fundamental role in the electrical priming effect observed there.

**Figure 2:**
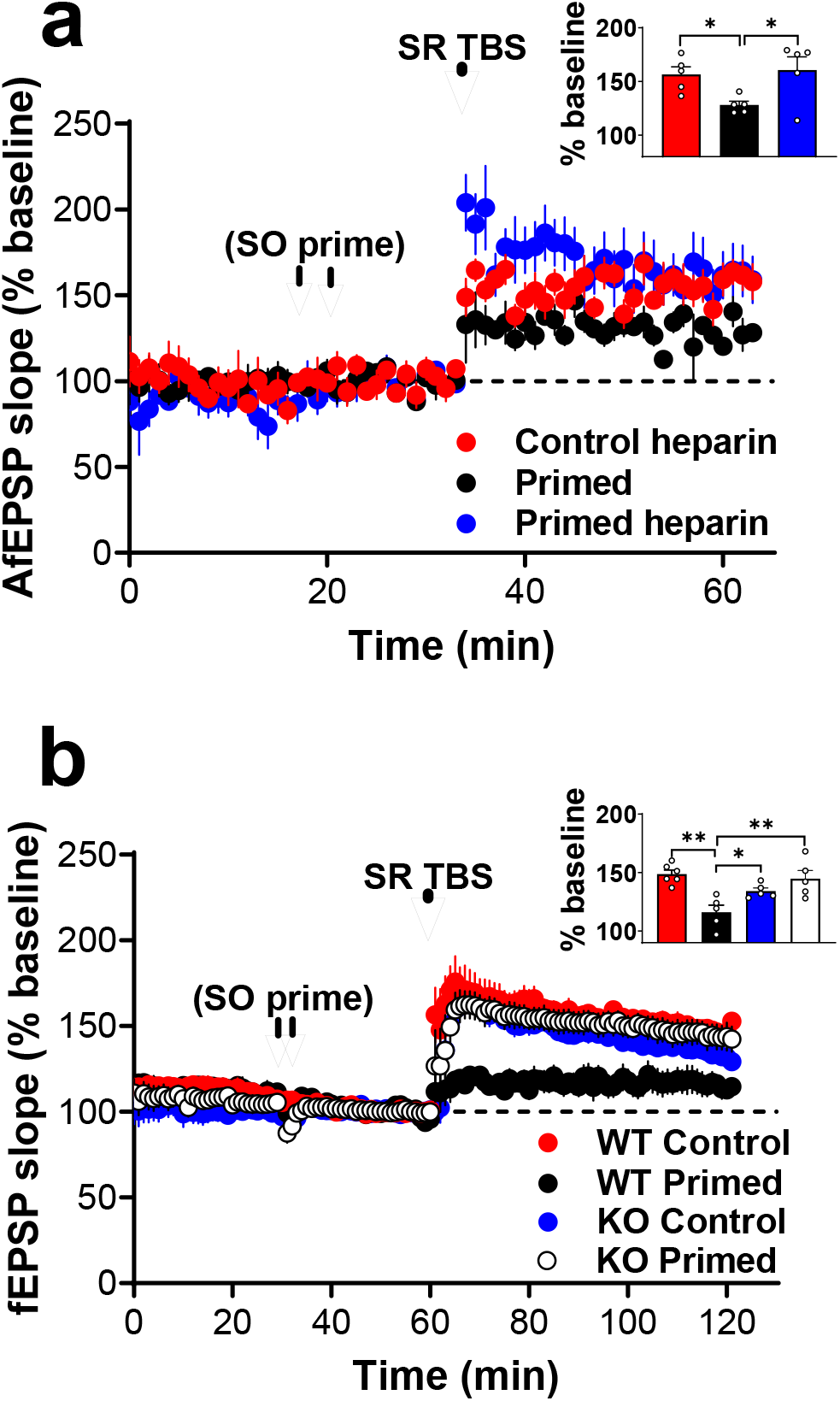
Astrocytic inositol triphosphate-gated stores are required for the metaplasticity effect. **a**, Dialysing single SR astrocytes with heparin blocked the metaplasticity effect (Control + Heparin (n=%): 157 ± 7%; Primed (n=5): 128 ± 4%; Primed + Heparin (n=5): 161 ± 12%; *F*_(2,12)_ *= 4.39, p* = 0.037). Inset: Bar chart of average AfEPSP LTP during the final 10 min of recording. **b**, The metaplasticity effect was absent in slices from *IP3R2KO* mice but not their wild type littermates (WT Control (n=5): 148 ± 4%; WT Primed (n=5): 116 ± 6%; KO Control (n=5): 134 ± 3%; KO Primed (n=5): 124 ± 7%; *F*_(3,17)_ = 8.553, *p* = 0.001). Inset: Barchart of average fEPSP LTP during the final 10 min of recording.

### 3.3 A_2B_Rs trigger astrocyte-dependent hippocampal metaplasticity

We previously reported that A_2B_R activation is both necessary and sufficient for triggering the metaplasticity effect (Jones et al., 2013). We demonstrated here that this A_2B_R-induced metaplasticity is also astrocyte-dependent, as brief bath-application of the A_2B_R agonist BAY-60 6583 inhibited LTP 30 min later at SR synapses near control non-clamped astrocytes, but not at synapses near calcium-clamped astrocytes (**Fig. 3a)**. We also tested whether the A_2B_R agonist could generate the metaplastic inhibition of LTP *in vivo*, using rats tested >4 weeks after chronically implanting electrodes into CA1 (a critical recovery period for avoiding aberrant release of cytokines from astrocytes following LTP induction *in vivo*: Jankowsky, Derrick, & Patterson, 2000). Under these strict conditions, brief intraventricular delivery of BAY-60 6583 30 min prior to LTP induction significantly attenuated SR LTP. This effect was blocked by co-administration of the A_2B_R antagonist MRS1754 (**Fig. 3b**). These data, together with the evidence that A_2B_R activation generates Ca^2+^ transients in astrocytes (Kawamura & Kawamura, 2011; Tanaka et al., 2021), provides further strong support for a role by astrocytes in mediating the metaplasticity effect.

**Figure 3:**
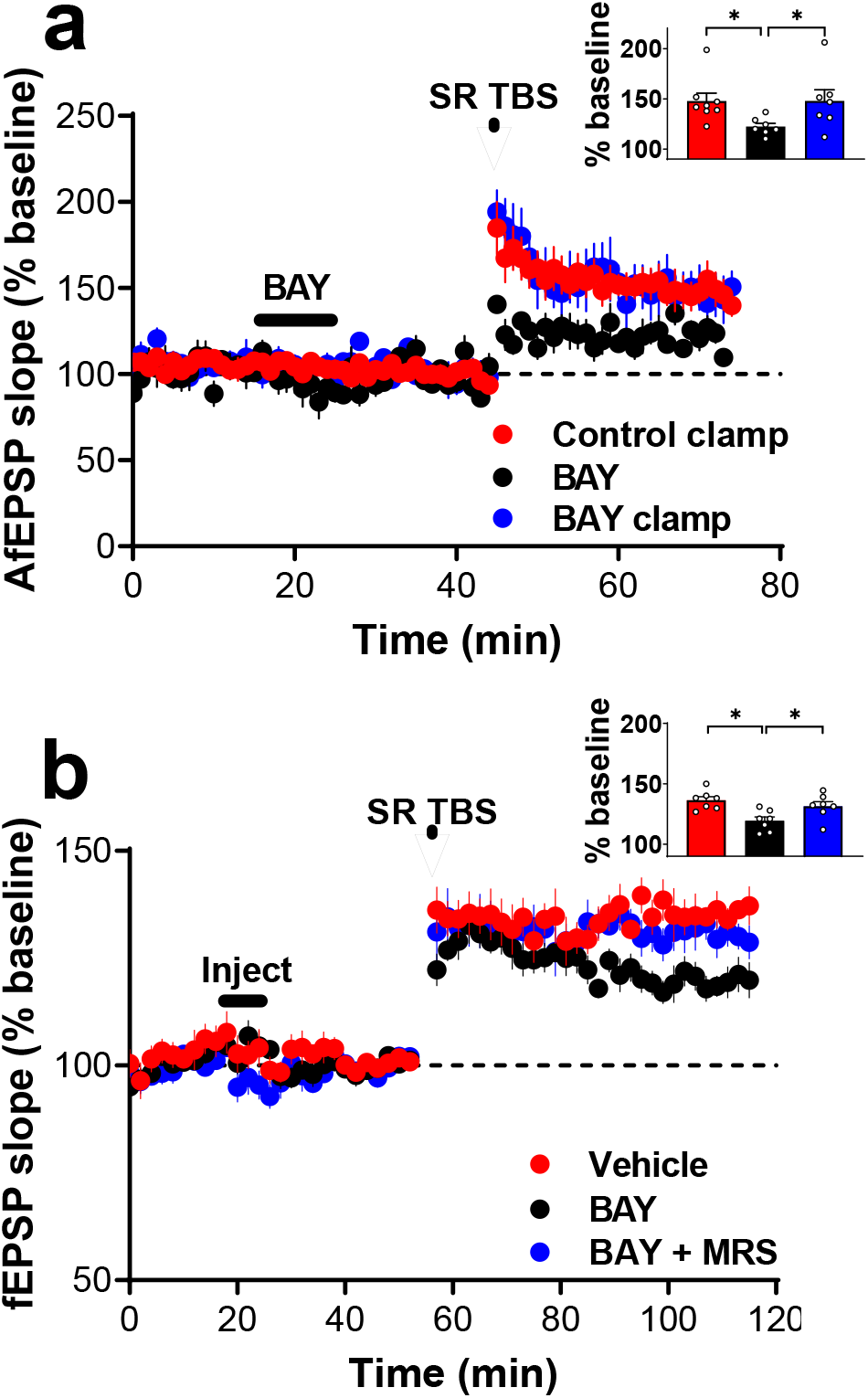
A_2B_Rs trigger astrocyte-dependent hippocampal metaplasticity. **a,** The astrocyte Ca^2+^ clamp blocked the inhibition of LTP by A_2B_R agonist priming (black bar) *in vitro* (Control + Clamp (n=8): 148 ± 8%; Primed (n=8): 122 ± 4%; Primed +Clamp (n=8): 148 ± 11%; *F*_(2,19)_ = 4.2, *p* = 0.032). Inset: Bar chart of average AfEPSP LTP during the final 10 min of recording. **b**, A_2B_R priming (black bar) also inhibited SR LTP *in vivo*, an effect blocked by MRS1754 (Vehicle control (n=7): 136 ± 3%; BAY-60 6583 (n=7): 119 ± 3%; BAY-60 6583 + MRS 1754 (n=7): 130 ± 3%; *F*_(2,18)_ = 7.18, *p* = 0.005). Inset: Bar chart of average fEPSP LTP during the final 10 min of recording.

### 3.4 Astrocyte-dependent metaplasticity via TNF and GluN2B-containing NMDA receptors

We have previously reported that the long-range metaplasticity effect requires the cytokine tumor necrosis factor (TNF) (Singh et al., 2019). TNF is a known gliotransmitter (Beattie et al., 2002; Canedo et al., 2021), and so we hypothesised that this cytokine may be the final astrocyte-to-neuron signal that ultimately inhibits LTP. Here we confirmed that bath administration of exogenous TNF protein (1.18 nM) for 10 min inhibits SR LTP induced 30 min later (**Fig. 4a)**. Contrary to our prediction, however, pharmacological priming with TNF also required astrocyte Ca^2+^, as the inhibition of LTP produced by this protocol was blocked at synapses near Ca^2+^-clamped astrocytes (**Fig. 4a)**. Thus, TNF acts upstream of astrocyte Ca^2+^ signaling and is not the final signal released onto neurons.

**Figure 4:**
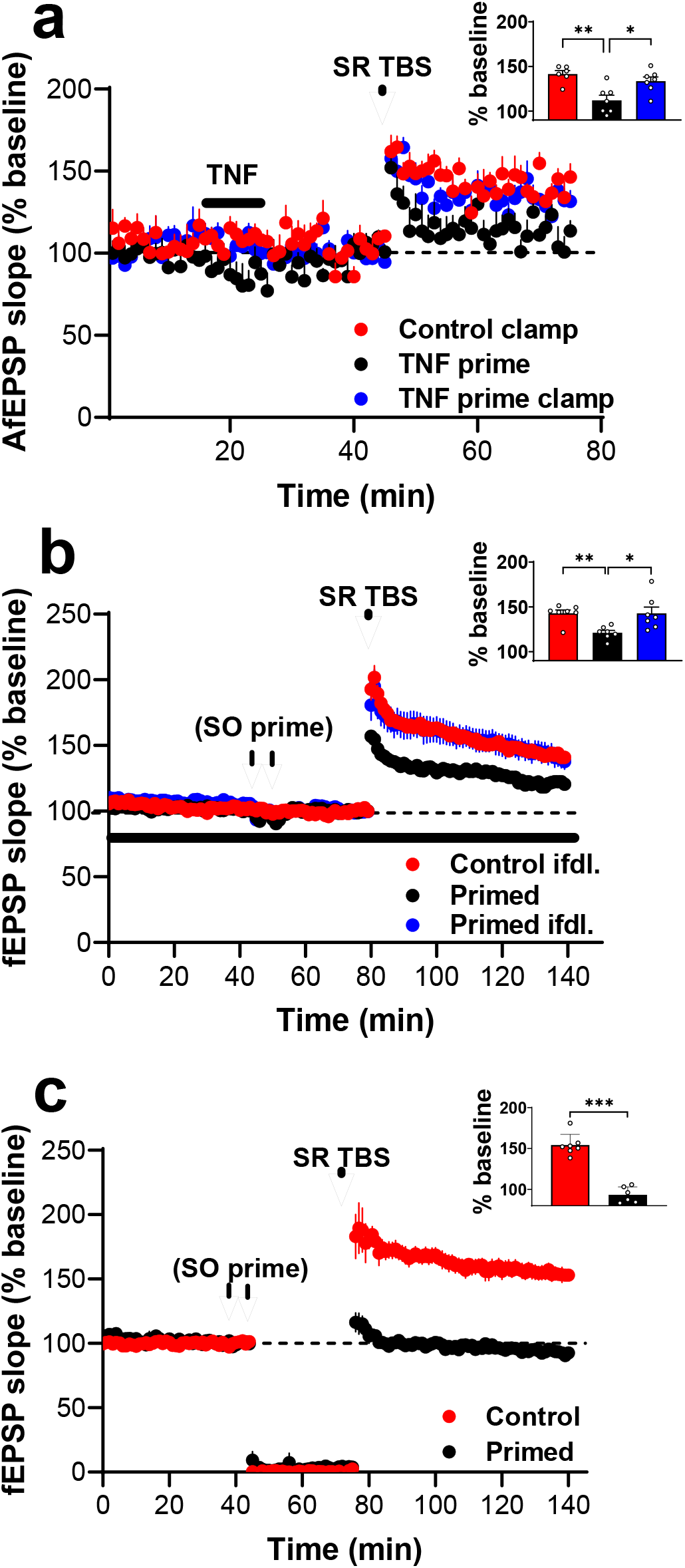
Astrocyte-dependent metaplasticity via TNF and GluN2B-containing NMDA receptors. **a**, TNF application inhibited LTP at SR synapses under control conditions, but not when astrocyte Ca^2+^ was clamped (Control clamp (n=7): 142 ± 7%; Primed (n=7): 112 ± 4%; Primed clamp (n=7): 134 ± 6%; *F*_(2,18)_ = 9.556, *p* = 0.002). Inset: Bar chart of average AfEPSP LTP during the final 10 min of recording. **b**, The metaplasticity was absent when Ifenprodil was perfused throughout the experiment (Control Ifenprodil (n=7): 143 ± 4%; Primed (n=7): 121 ± 2%; Primed Ifenprodil (n=7): 143 ± 3%; *F*_(2,18)_ = 6.629, *p* = 0.007). Inset: Bar chart of average fEPSP LTP during the final 10 min of recording. **c**, The metaplasticity was evident even when synaptic stimulation was stopped between priming and later LTP induction (Control (n=7): 154 ± 5%; Primed (n=6): 93 ± 4%; *t*_(1,13)_ = 9.32, *p* < 0.001.

It is known that TNF can trigger glutamate release from astrocytes, which acts predominantly on neuronal NMDA receptors containing the GluN2B subunit (Habbas et al., 2015; Santello, Bezzi, & Volterra, 2011). Further, there is evidence that TNF-mediated glutamate release from astrocytes is dependent on IP_3_R-mediated Ca^2+^release (Canedo et al., 2021) and the additional release of ATP (Nikolic et al., 2018), as per our effect (Jones et al., 2013). This glutamate release is ongoing, outlasting the duration of TNF application (Canedo et al., 2021), which is reminiscent of the ongoing Ca^2+^release in astrocytes triggered by brief A_2B_R activation (Kawamura & Kawamura, 2011). On the other hand, we know that NMDARs are not involved in the present effect when blocked before, during and shortly after the priming stimulation (**Fig. 1a)**. Taken together, these findings led us to ask whether astrocytes might remain spontaneously active post-priming, and release glutamate in an ongoing manner that outlasts the washout period of glutamatergic antagonists in our earlier experiment (**Fig. 1a)**. Thus, we undertook two sets of experiments to determine whether ongoing spontaneous transmission onto GluN2B-containing receptors could be required for the metaplasticity effect. First, we conducted our standard electrical priming experiments and pharmacologically inhibited GluN2B-containing NMDARs throughout the experiment using ifenprodil (3 µM), a paradigm that does not block basal levels of LTP in CA1 (Papouin et al., 2012; Yang et al., 2017). When ifenprodil was present for the duration of the experiment (i.e., not only during priming), priming had no significant effect on later LTP (**Fig. 4b**). The metaplasticity therefore requires ongoing activation of GluN2B-containing receptors after priming and close to the time of LTP induction.

We then probed whether the relevant glutamate could come from our synaptic test-pulse stimulation instead of astrocytes, by ceasing stimulation between priming and later LTP induction. Even under these conditions, later LTP was induced normally in non-primed slices but the priming stimulation readily inhibited it (**Fig. 4C**). Indeed, whereas some residual LTP is usually evident in SR even after priming, LTP was completely absent in primed slices in these experiments, suggesting that ongoing stimulation may in fact partly counteract the effects of priming. The relevant source of glutamate required for the effects of priming is therefore likely non-synaptic.

## 4. Discussion

Our findings demonstrate that astrocytes play a fundamental role in sensing and responding to activity in the hippocampal network by generating a long-range inhibition of subsequent LTP. This mode of metaplasticity occurs independently of conventional excitatory or inhibitory synaptic transmission, relying instead on an astrocyte-mediated cascade including A_2B_R-, IP_3_R2-, TNF-and GluN2B-dependent mechanisms.

Our results provide important advances to the understanding Ca^2+^-dependent neuron-astrocyte bi-directional communication. While it has been appreciated for some time that astrocytic Ca^2+^ is an essential requirement for NMDAR-dependent LTP induction (Henneberger et al., 2010), we now show that IP_3_R-dependent Ca^2+^ stores in astrocytes can also serve as a mechanism for generating the opposite effect, i.e. an inhibition of LTP. On the other hand, Ca^2+^ influx through TRP1A channels is responsible for driving the release of serine from astrocytes (Shigetomi, Jackson-Weaver, Huckstepp, O’Dell, & Khakh, 2013). Thus, it appears that different sources of astrocytic Ca^2+^ can exert opposing effects on LTP at CA1 SR synapses. Of relevance to this, it was recently reported that memory consolidation is impaired in IP3R2KO mice (Pinto-Duarte, Roberts, Ouyang, & Sejnowski, 2019), an effect ascribed to deficient long-term depression (LTD) at SR synapses in these animals. Our SO priming protocol actually *enhances* LTD at SR synapses, in addition to inhibiting LTP (Hulme et al., 2012). The metaplastic state may therefore be an astrocyte-mediated mechanism that primes long-term memory storage through the dual actions of LTP suppression and LTD enhancement (believed to be two key factors in consolidation (Vyazovskiy, Cirelli, Pfister-Genskow, Faraguna, & Tononi, 2008)).

A second important consequence of our results is the identification of a functional consequence to long-range astrocytic signaling. Astrocytes are known to signal over considerable distances via intercellular Ca^2+^ “waves” (Kuga, Sasaki, Takahara, Matsuki, & Ikegaya, 2011), but the downstream effects of these waves are seldom reported. Indeed, while early investigations sought to establish a role for such events in neurovascular coupling and cortical spreading depression, it is now generally accepted that astrocyte Ca^2+^ elevations are secondary to both effects (Bonder & McCarthy, 2014; Nizar et al., 2013; Peters, Schipke, Hashimoto, & Kettenmann, 2003; Wu et al., 2018). While we have previously reported that SO priming elicits Ca^2+^ elevations in astrocytes that spread as far as SR (Hulme et al., 2014), it is not until now that we have been able to demonstrate a causal link between astrocytic Ca^2+^ mechanisms and the induction of the metaplastic state in SR. To our knowledge, this is the first report of a functional consequence of long-distance astrocytic signalling. As astrocyte Ca^2+^ waves spread across many cells in awake, mobile animals (Ding et al., 2013; Dombeck, Khabbaz, Collman, Adelman, & Tank, 2007; Nimmerjahn, Mukamel, & Schnitzer, 2009; Paukert et al., 2014), we suggest the astrocyte-mediated spread of metaplastic states as a plausible, behaviourally relevant mechanism for regulating neural networks over broad spatiotemporal scales.

A_2B_Rs generate enduring (>20 min) and spatially widespread (>300 µm) release of Ca^2+^within the hippocampal astrocytic syncytium (Kawamura & Kawamura, 2011). In keeping with these spatiotemporal scales, we had previously demonstrated that pressure ejection of BAY-60 6583 into SO generates a metaplastic state that spreads to SR (Jones et al., 2013). We now provide the crucial link between these findings, demonstrating that BAY-606583 priming acts via astrocytes, as hypothesised. We also show that TNF contributes to the metaplasticity in an astrocyte-dependent manner, and that GluN2B-containing NMDARs are required. The involvement of these two molecules is important not only because they are functionally linked (Habbas et al., 2015) but also because both TNF and GluN2B-NMDARs have known roles as inhibitors of LTP (Butler, O’Connor, & Moynagh, 2004; S. Li et al., 2011; Olsen & Sheng, 2012; Wang, Wu, Rowan, & Anwyl, 2005).

But how do the many disparate molecules in our model connect? One straight-forward explanation is that activation of A_2B_Rs by priming stimulation triggers the widespread release of TNF, which in turn triggers the release of glutamate from astrocytes onto nearby neurons. Yet, this interpretation is problematic as, while A_2B_Rs are known to trigger release of cytokines (Moidunny et al., 2012; Vazquez, Clement, Sommer, Schulz, & Van Calker, 2008), there is no evidence to date that this includes TNF. An alternative explanation is that TNF lies upstream from A_2B_R activation. TNF treatment causes phosphorylation of A_2B_Rs on human astroglia (Trincavelli, Tonazzini, Montali, Abbracchio, & Martini, 2008), and TNF is an upstream, permissive factor for ATP-driven Ca^2+^-dependent glutamate release from astrocytes (Nikolic et al., 2018; Santello et al., 2011). It is therefore possible that TNF directly or indirectly leads to A_2B_R-dependent Ca^2+^ elevations.

A final consideration is the significance of our results for how neural network activity is stabilised. Computational models of learning, such as the BCM model, often include regulatory mechanisms that maintain activity or plasticity within optimal bounds, but such models are typically based on mechanisms that are intrinsic to the postsynaptic neuron. Our results present a departure from this neuron-centric view. Astrocytes inhabit discreet, non-overlapping territories (Bushong, Martone, Jones, & Ellisman, 2002) while at the same time forming a vast, interconnected network. Astrocytes are therefore well-placed to influence synapses over a range of spatial scales (much broader than a single neuron). Astrocyte-mediated metaplasticity as seen in our model may reflect a transient response to behavioural experience, or a chronic response under pathological conditions (Singh et al., 2019), where it is favourable to dampen future LTP that could in lead to further instability in the network.

## Acknowledgements

IP3R2KO mice were generously provided by Dr Ju Chen and the University of California, San Diego.

## Supplementary information

**Supplementary Figure 1.**
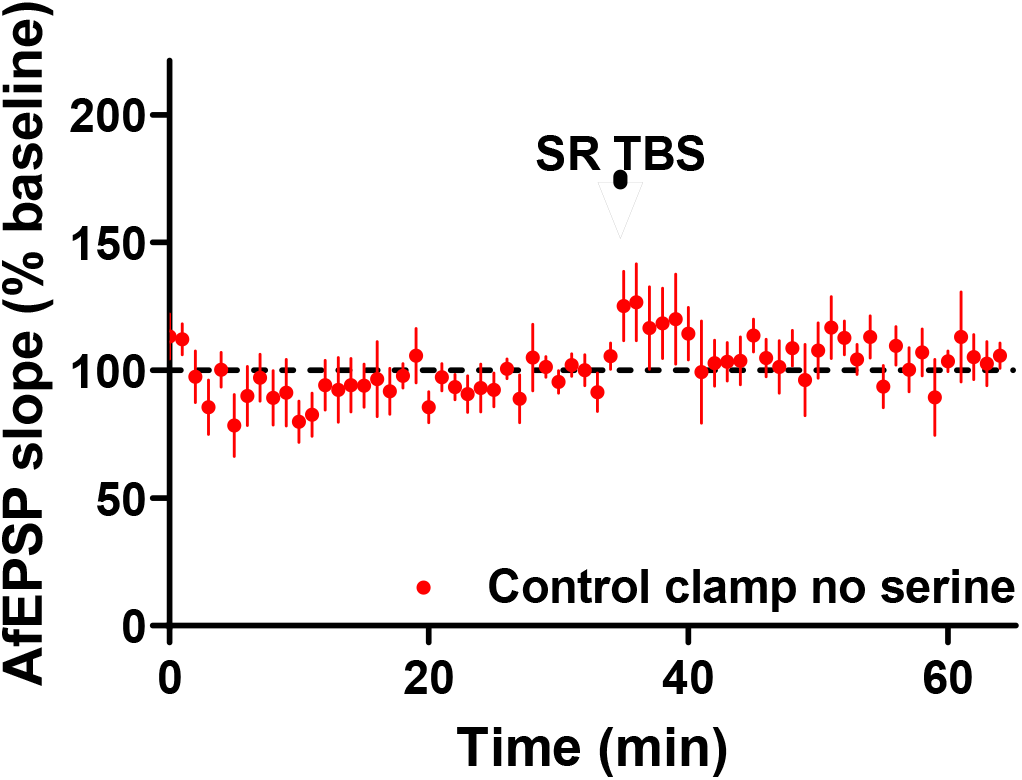
LTP was not elicited near astrocytes whose cytosolic Ca^2+^ was clamped with EGTA and D-serine was not present in the ACSF (Control clamp, no serine (n=7), pre vs. post TBS: 100 vs. 104 ± 6%, paired *t*_(6)_ = 2.45, *p* = 0.52). The same TBS paradigm in the presence of bath-applied D-serine induced a substantial LTP (cf. **Fig. 1b**).

